# Local Control of Intracellular Microtubule Dynamics by End Binding Protein 1 (EB1) Photo-Dissociation

**DOI:** 10.1101/099598

**Authors:** Jeffrey van Haren, Andreas Ettinger, Hui Wang, Klaus M. Hahn, Torsten Wittmann

## Abstract

Dynamic remodelling of the microtubule cytoskeleton and local interactions with intracellular targets are central to many polarized cell biological processes, an idea first formalized as search-and-capture hypothesis three decades ago^1^. However, because of the rapid timescale of microtubule polymerization dynamics, it is difficult to directly ask how, when and where dynamic microtubules participate in specific biological processes. Here, we employ a blue light-sensitive interaction with the oat phototropin LOV2 domain^2^ to generate a photo-inactivated variant of the microtubule end-binding protein EB1, a small adaptor that is central to the interaction of functionally and structurally diverse proteins with growing microtubule ends^3,4^, that can replace endogenous EB1 function. Acute and reversible blue light-mediated n-EB1 photo-dissociation allows spatially and temporally precise control of intracellular microtubule polymerization dynamics. In addition to demonstrating that neither the GTP cap nor the MT polymerase CKAP5 are sufficient to sustain persistent MT polymerization at physiological growth rates, our data illustrate accurate subcellular control of a freely diffusible, cytoplasmic protein at the second and micrometer scale. This novel design may serve as a template for precise control of many other intracellular protein activities.

---

Control of microtubule (MT) polymerization dynamics and interactions of growing MTs with other intracellular components are mediated by a heterogeneous class of proteins commonly referred to as +TIPs^3,4^. Most if not all +TIP interactions with growing MT ends require end-binding proteins (EBs). However, different +TIPs can have antagonistic activities. For example, interactions with EBs can recruit enzymes to growing MT ends promoting either MT polymerization^5^ or depolymerization^6^, and it is not known how these opposing activities are spatially and temporally balanced in cells. It also remains unclear how MT dynamics and the +TIP network of proteins contribute to complex cell and tissue morphogenesis, which represents a significant gap in our understanding of physiological MT function^7^. Because genetic methods are orders of magnitude too slow to dissect highly dynamic +TIP functions in cells with sufficient spatial and temporal resolution, we targeted the +TIP adaptor activity of EB1 using a new ‘opto-cell biology’ approach.

EB1 is a small dimer that consists of two functional domains connected by an intrinsically disordered linker. The N-terminal CH domain recognizes the guanosine nucleotide state of growing MT ends^8,9^ while the C-terminal EBH domain is required to recruit +TIPs to growing MT plus ends^10^. We predicted that controlling the connection of these two functional halves using a light-controlled protein interaction pair would allow us to modulate +TIP functions in cells with high accuracy. While most light-sensitive protein interactions are induced by light^11^, the recently described Protein A Z-domain-derived affibody, Zdk1^2^, functions the opposite way and binds the oat phototropin LOV2 domain with high affinity in the dark, but dissociates from LOV2 in blue light (Fig. 1a). In addition, the LOV2/Zdk1 pair is highly soluble, does not require exogenous co-factors in mammalian cells, and its molecular weight is less than that of GFP, and therefore should interfere minimally with protein function.

**Figure 1.**
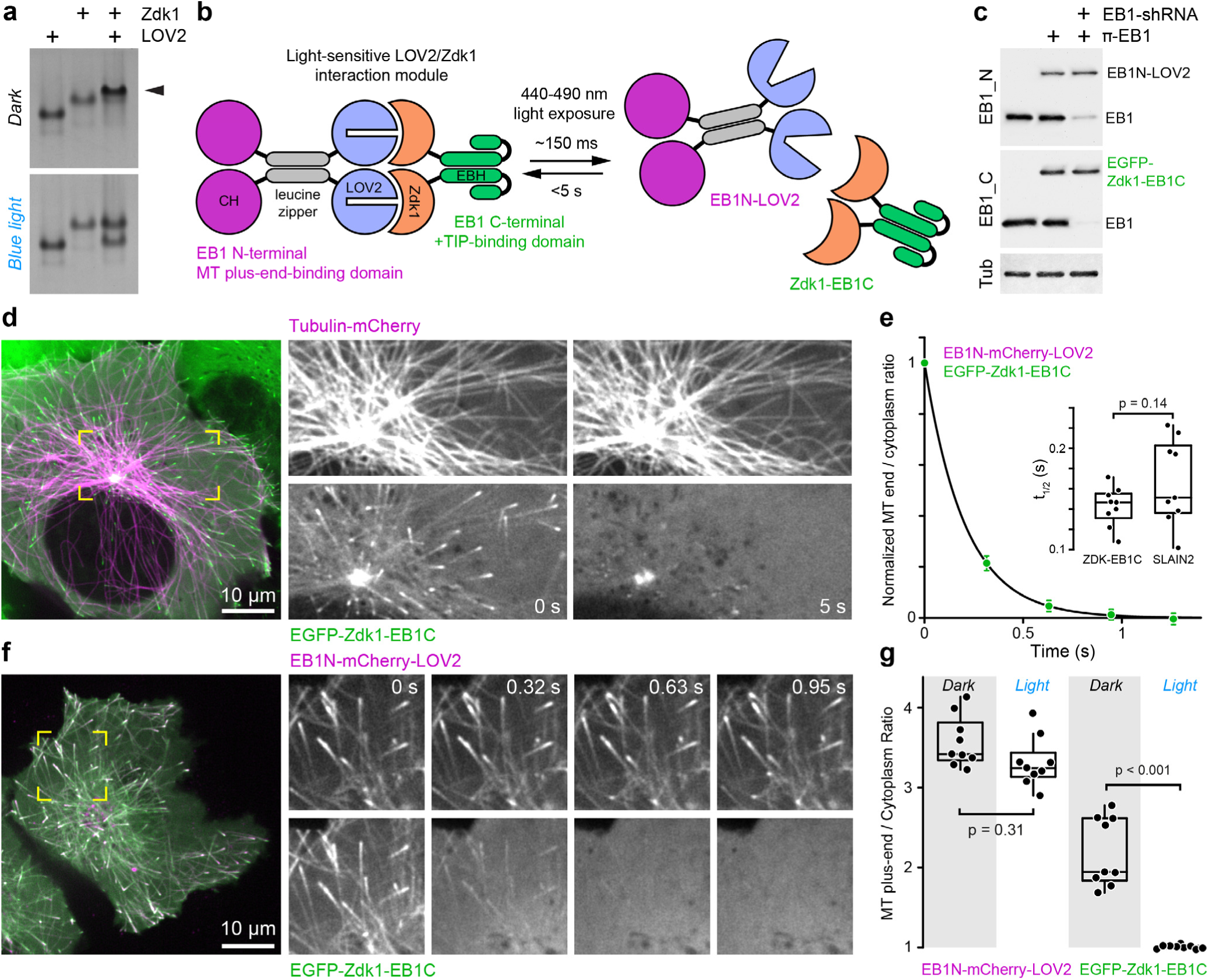
A light-sensitive EB1 variant can replace endogenous EB1 function. **(a)** Interaction of purified LOV2 domain and Zdk1 peptide analysed by native PAGE. Blue light results in dissociation of the LOV2/Zdk1 complex (arrow) that is upshifted compared to LOV2 or Zdk1. **(b)**Schematic of the photo-inactivated π-EB1 design resulting in reversible dissociation of the N-terminal MT-binding and the C-terminal +TIP adapter domains upon blue light exposure. **(c)** Analysis of EB1 and π-EB1 expression levels in H1299 cell lines in which π-EB1 constructs were stably expressed and endogenous EB1 depleted by shRNA. Immunoblots were probed with antibodies specific to either the C- or N-terminus of EB1. **(d)** Cell expressing Tubulin-mCherry and EGFP-tagged Zdk1-EB1C before and after 5 s of exposure to 488 nm blue light resulting in dissociation of Zdk1-EB1C from growing MT plus ends. **(e)** Analysis of the blue light-induced EGFP-Zdk1-EB1C dissociation rate from MT ends. n = 9 cells. Error bars are 95% confidence intervals. Solid line is a fit with an exponential decay. Inset shows the comparison of dissociation half-lives of Zdk1-EB1C and SLAIN2, a +TIP that depends on EB1 for MT end association. **(f)** Cell expressing both halves of π-EB1 fluorescently tagged showing that EB1N-LOV2 remains on growing MT ends after blue light exposure. Time stamps indicate duration of blue light exposure. Dual-wavelength images were acquired simultaneously using an emission image splitter. **(g)** Analysis of the amount of the two π-EB1 halves bound to MT ends before and during 1 s of blue light exposure. In e and g, each symbol represents the average of 6-8 measurements from one cell. p-values were determined by Tukey-Kramer HSD test.

To develop a light-sensitive EB1 variant, we thus inserted the LOV2/Zdk1 module between the N-terminal and C-terminal halves of EB1 with the expectation that light-induced dissociation of these constructs would disrupt EB-mediated +TIP interactions with growing MT ends, without interfering with EBs ability to bind MTs. Because dimerization is required for efficient EB plus-end-tracking^12,13^, we further inserted a GCN4 leucine zipper (LZ) that is structurally similar to the EBH domain between the CH and LOV2 domains to retain dimerization and plus-end-tracking of the N-terminal half by itself. We refer to these photo-inactivated π-EB1 constructs as EB1N-LOV2 and Zdk1-EB1C (Fig. 1b). Bacterially expressed GST-EB1N-LOV2 bound 6xHis-Zdk1-EB1C and precipitated both SxIP motif (CLASP2) and CAP-Gly motif (p150_Glued_) containing +TIPs from cell lysates indicating that the two π-EB1 halves interact and that Zdk1-EB1C is functional in binding known classes of +TIPs (Supplementary Fig. 1a). To directly visualize π-EB! dynamics in live cells and because tagging EB1 at either the N- or C-termini interferes with EB1 function^14^, we inserted red or green FP tags N-terminal of the Zdk1 peptide. In these initial experiments to test the light response of the EB1N-LOV2 / Zdk1-EB1C interaction we photoactivated the LOV2 domain simply by turning on the 488 nm acquisition channel, which represents saturating blue light exposure. EGFP-Zdk1-EB1C was efficiently recruited to MT ends by non-tagged EB1N-LOV2, and dissociated from MT ends in response to blue light exposure with a half-life below 200 ms (Fig. 1d, e), which is close to the diffusion-limited turnover time of EB1 molecules on MT ends^15,16^. Of note, Zdk1-EB1C remained enriched near the centrosomes that contain SxIP-motif +TIPs independent of growing MT ends^17^. Because the LOV2 C-terminus is required for Zdk1-binding, we further inserted a different FP tag between EB1N and LOV2 demonstrating that at short time scales plus-end-tracking of EB1N-LOV2 was not significantly reduced by π-EB1 photo dissociation (Fig. 1f, g). These data demonstrate that both halves of π-EB1 are recruited to MT ends efficiently in the dark, and that blue light induces extraordinarily rapid π-EB 1 photo-dissociation.

In order to test if π-EB1 was functional, we generated cell lines stably expressing π-EB1 constructs in which we subsequently depleted endogenous EB1 by lentivirus-mediated shRNA (Fig. 1c). We chose H1299 non-small lung cancer cells because these cells express around twenty times more EB1 than EB3^18^. Consequently, unlike anti-EB1 antibodies, anti-EB3 antibodies only labelled MT ends in cells also expressing EB3-mCherry (Supplementary Fig 1c). EB3-mCherry also partially dissociated from MT ends upon π-EB1 photo-dissociation, likely reflecting some degree of heterodimerization with π-EB 1 and supporting a dominant effect of π-EB1 inactivation even in the presence of small amounts of endogenous EBs (Supplementary Fig. 2a)^13,19^. Although the π-EB1 H1299 cell lines expressed the EB1N-LOV2 and Zdk1-EB1C constructs at a slightly lower level compared with endogenous EB1 in control cells (Fig. 1c), these cells that are used in subsequent functional experiments were viable with no obvious phenotype, indicating that π-EB1 can replace endogenous EB1 function.

In addition, all EB-dependent +TIPs tested tracked MT plus ends in non-illuminated cells, but disappeared from MT ends with similarly fast kinetics as Zdk1-EB1C during blue light exposure (Figs. 2a and 1e) confirming that MT end association of MCAK, CLASP2, and SLAIN2 critically depends on interactions with the C-terminal domain of EB1. +TIP dissociation was further reversible over multiple cycles of light exposure (Fig. 2b; Supplementary Video 1) indicating that π-EB1 photo-dissociation rapidly and reversibly disrupts the MT plus end +TIP complex. +TIPs only dissociated from growing MT ends and not from other binding sites; e.g. CLASP2 remained at the Golgi apparatus^20^ (Supplementary Fig. 2b), demonstrating that +TIP dissociation from MT plus ends is specific to the light-sensitive dissociation of the two π-EB1 halves.

**Figure 2.**
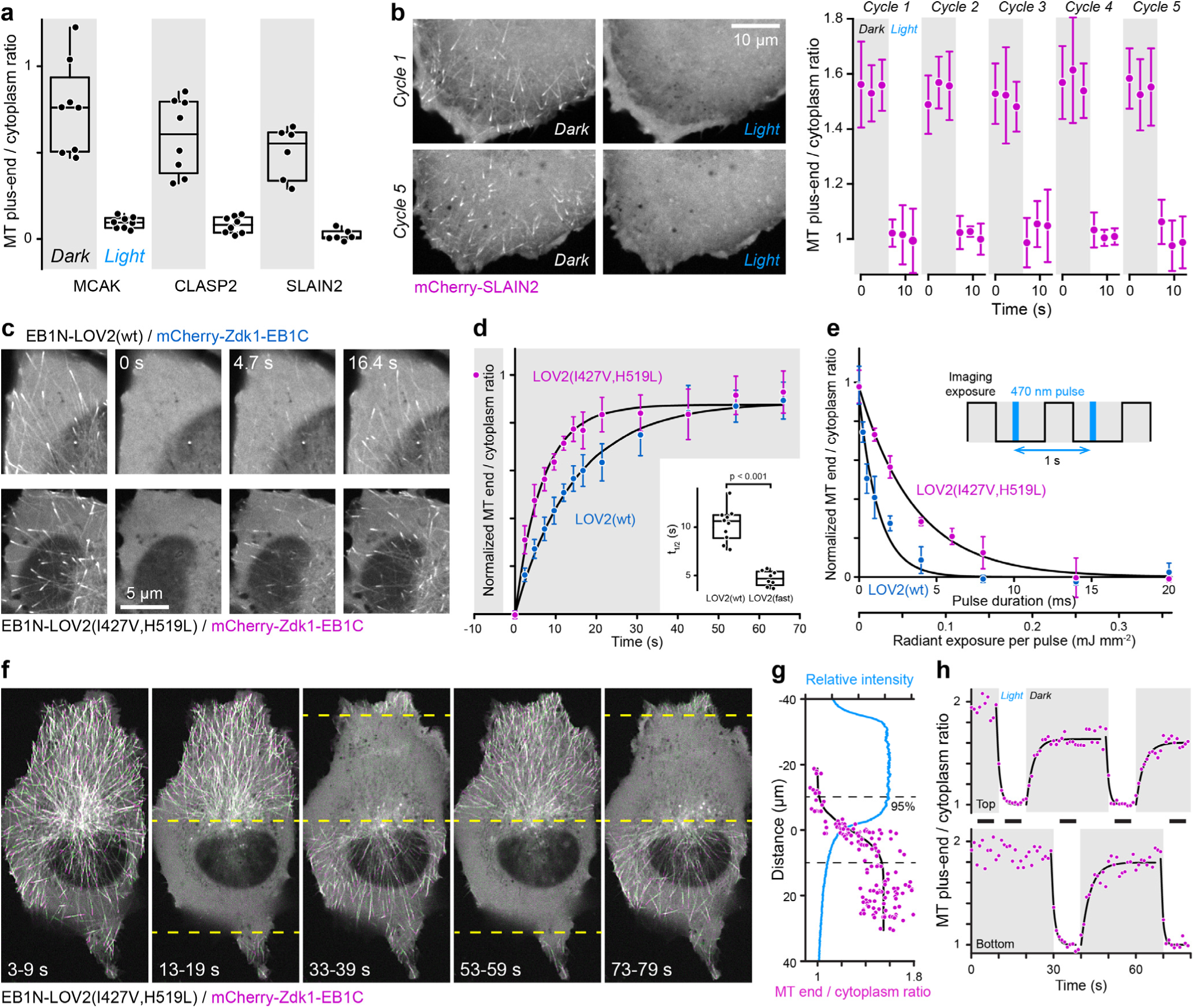
Spatially and temporally reversible photo-dissociation of +TIP complexes. **(a)** Analysis of the amount of the indicated EB1-dependent +TIPs bound to MT ends before and during 1 s of blue light exposure in π-EB1 / EB1 shRNA cells. Each symbol represents the average of 10 measurements per cell. **(b)**Reversible blue light-induced disruption of mCherry-SLAIN2 binding to MT ends in π-EB1 / EB1 shRNA cells. Graph shows mCherry-SLAIN2 MT end association over multiple cycles of blue light exposure. n = 6 MT ends. Error bars show standard deviation. **(c)** Comparison of Zdk1-EB1C recovery on MT plus ends in cells expressing π-EB1 containing either wild-type LOV2 or the LOV2(I427V,H519L) variant. Time stamps indicate time elapsed after blue light exposure was terminated. **(d)** Analysis of recovery rates of π-EB1 containing the indicated LOV2 variants. n = 8-10 cells. Error bars are 95% confidence intervals. Solid lines are exponential curve fits. Inset shows the comparison of recovery half-lives. **(e)** Analysis of Zdk1-EB1C MT plus end association in π-EB1 constructs with the indicated LOV2 variants as a function of blue light radiant exposure. Total blue light dose was varied by pulse width modulation as illustrated in the inset. Measurements were taken at steady state after 40 s of pulsed blue light exposure showing tuning of π-EB1 photo-dissociation at intermediate exposure levels. n = 3-4 cells per pulse length. Error bars are 95% confidence intervals. Solid lines are exponential curve fits. **(f)** Patterned blue light exposure within the regions indicated by dashed lines shows high spatial and temporal accuracy of π-EB1 photo-dissociation. In this example, the blue light pattern is switched every 10 s. Images are projections of the indicated time periods showing sequential time points in alternating green and purple. These time periods are also indicated as black bars in h. **(g)** Analysis of Zdk1-EB1C MT plus end association at the boundary of the blue light pattern corresponding to the 33-39 s period showing the best achievable spatial control. Solid line is a fit with a cumulative normal distribution. Dashed lines demarcate the 95% switch (i.e. +/- 2 σ) in MT end association calculated from the fit. (h) Analysis of Zdk1-EB1C MT plus end association as function of time in the top and bottom halves of the cell shown in f. Solid lines are exponential curve fits. Shaded areas indicate time periods without blue light exposure.

Because π-EB1 likely diffuses freely through the cytoplasm^16^, spatial control inside cells requires rapid return of photoactivated π-EB1 molecules to the dark state to prevent diffusion of activated molecules throughout the cell. Compared with wild-type LOV2, re-association of π-EB1 containing the previously described I427V LOV2 variant^2,21^ occurred approximately twice as fast with a half-life of less than 5 s at saturating blue light levels (Fig. 2c, d). To minimize blue light-induced phototoxicity through potential FMN decomposition and singlet oxygen generation^22^, we next determined the minimum blue light dosage required for sustained π-EB1 photo-dissociation. Because most light sources cannot be reliably modulated over a sufficiently wide range of intensities, we instead used pulse width modulation to achieve specific radiant exposures by briefly flashing a 470 nm LED at a given irradiance level (∼18 mW mm^-2^) at a frequency of 1 Hz between camera exposures (Fig. 2e). Compared with wild-type LOV2, an approximately three times increased blue light exposure was needed to reach steady state at which half of π-EB1 with the faster LOV2 domain variant dissociated from MT ends, indicating that faster LOV2 dark recovery comes at a cost of increased blue light exposure needed for photoactivation. Importantly, however, for both LOV2 variants the required radiant exposure to achieve complete π-EB1 photo-dissociation is several orders of magnitude below light doses encountered in fluorescence microscopy, which can easily exceed 100 mJ mm^-2^ per exposure at typical irradiance levels^23^, suggesting that the photodamage associated with π-EB1 photoactivation should be negligible.

Next, we used a mask generated with a digital micromirror device (DMD) to test if LOV2 photodynamics are sufficient to achieve local π-EB1 inactivation at optimized exposure levels. Using the faster LOV2 variant, 20 ms 470 nm light pulses of ˜1 mJ mm^-2^ every second (i.e. a 2% duty cycle) were sufficient to achieve intracellular gradients with 95% difference in π-EB1 photo-dissociation over a width of ˜20 μm (Fig. 2f, g), and to reversibly switch π-EB1 photo-dissociation between different intracellular regions (Fig. 2h; Supplementary Video 2). To our surprise, wild-type LOV2 achieved a boundary that was almost as steep at an lower blue light exposure (10 ms, 1% duty cycle). In both cases, the boundary steepness was similar to the edge steepness of the illumination pattern (Fig. 2g), which we found to be significantly shallower than what could be theoretically expected from the optical resolution limit. This indicates that in our system boundary steepness is mostly determined by focus and scattering of the illumination pattern edge, which is a current technical limit, and less by biophysical properties such as diffusion of photo-dissociated π-EB1 molecules out of the blue light-exposed region. Because of the lower light dose required to activate wild-type LOV2, we therefore continued to use the wild-type LOV2 πEB1 for subsequent experiments.

Many +TIPs that are recruited to MT ends through interactions with EB1 directly or indirectly influence MT polymerization dynamics, but it is not understood how potentially antagonistic +TIP activities are integrated in cells and spatially controlled in real time^3^. Although previous siRNA depletion of EB proteins demonstrated defects in MT polymerization dynamics^13,24^, such experiments are confounded by indirect and adaptive mechanisms in genetically depleted cells and cannot directly address specific protein actions in real time. We therefore tested how π-EB1 photo-dissociation acutely altered MT dynamics and organization. Computational tracking of EB1N-mCherry-LOV2 that remains on growing MT ends upon π-EB1 photo-dissociation revealed an overall attenuation of MT growth and a reduction in the number of growing MT ends within 30 s of blue light exposure (Fig. 3a, b; Supplementary Video 3). Further analysis of growth rate distributions indicated that π-EB1 photodissociation specifically reduced the growth rate of a fast growing MT population (Fig. 3c) that likely represent persistently growing MTs in the cell interior^25^. Importantly, these changes were reversible, and MT dynamics nearly completely recovered within minutes of terminating blue light exposure (Fig. 3b). Moreover, the fast photodynamics of π-EB1 allowed for locally restricted control of MT growth dynamics (Fig. 3d; Supplementary Video 4). Analysis at higher temporal resolution further showed that individual MTs can continue fast growth for a short period of time before slowing down, becoming more dynamic and transitioning to depolymerisation (Fig. 3e) and that the growth rate of the MT population decayed with a half-life of ˜5 seconds following blue light exposure (Fig. 3f). Because this is an order of magnitude slower than π-EB1 photo-dissociation (Fig. 1e), it must reflect an intrinsic MT property and is in good agreement with the average life-time of the cap of high affinity EB1 binding sites determined *in vitro*^26^. This suggests that the GTP-like MT end cap can temporarily maintain a MT end structure permissive of fast MT growth. However, in contrast to *in vitro* where the MT end cap is thought to sufficiently protect slow polymerizing MTs^27^, our data show that in cells it cannot maintain continued persistent MT growth, which is ˜10 faster and occurs at limiting free tubulin concentrations. Similarly, the N-terminal MT-binding part of π-EB1 by itself, and thus structural changes induced by EB1 binding^9^, are insufficient to maintain persistent MT growth. Instead, direct EB1 effects on MT dynamics in cells are outweighed by the activities of the EB1- recruited +TIP complex.

**Figure 3.**
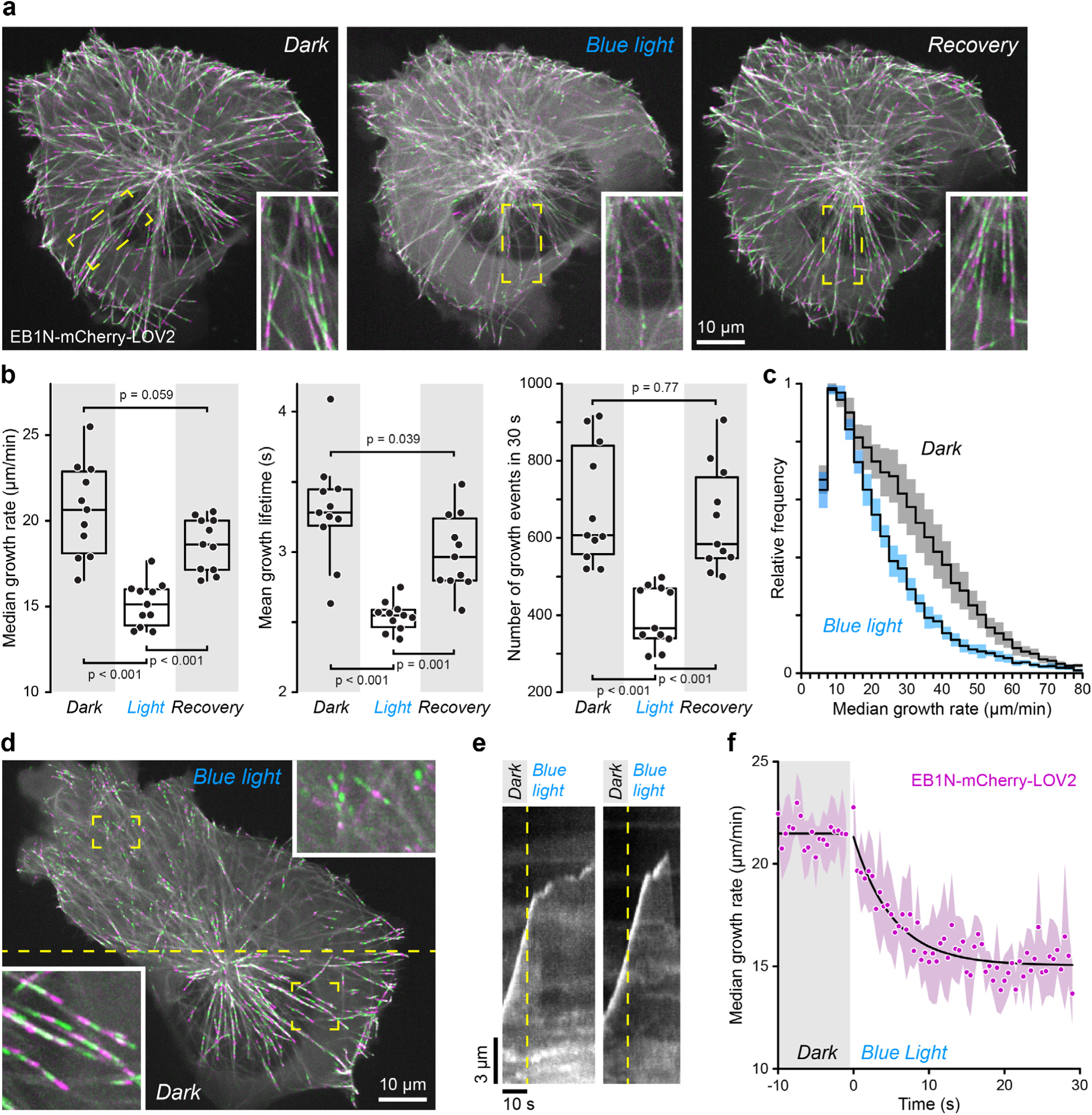
Attenuation of MT growth by π-EB1 photo-dissociation. **(a)** Projections of EB1N-mCherry-LOV2 labelled MT plus ends in 20 s temporal windows before, after 30 s during blue light exposure, and after 3 min recovery in the dark. Insets show the indicated regions at higher magnification. MT growth tracks appear in alternating colours. **(b)**Quantification of MT polymerization dynamics by tracking EB1N-mCherry-LOV2 labelled MT ends. Growth rates are frame-to-frame measurements from images acquired at 0.5 s intervals. Each symbol represents one cell. p-values were determined by Tukey-Kramer HSD test. **(c)** Comparison of the frame-to-frame MT growth rate distribution in the dark and during blue light exposure demonstrating a specific loss of fast growth events as a result of π-EB1 photo-dissociation. n = 11 cells. Shaded areas indicate 95% confidence intervals. **(d)** Local inhibition of MT growth by patterned blue light exposure. Only the top half of the cell shown was exposed to blue light pulses between image acquisitions. **(e)** Example kymographs illustrating the response of rapidly growing MT ends to π-EB1 photo-dissociation. **(f)** Quantitative analysis of the intracellular MT population growth rate response as a function of time after π-EB1 photo-dissociation. Solid lines are linear (before) and exponential fits (during blue light exposure). n = 5 cells. Shaded area indicates the 95% confidence interval.

These data also demonstrate that EB1-mediated recruitment of MT-polymerizing activities is dominant in interphase cells. Because SLAIN2, a +TIP that directly binds to and has been proposed to recruit the MT polymerase CKAP5^5^, was rapidly lost from MT ends following π-EB1 photo-dissociation (Fig. 1e; Fig. 2a, b) we next analysed the dynamics of CKAP5 in response to π-EB1 photo-dissociation. Although CKAP5-mKate2 dots were only visible by TIRF microscopy^28^ and difficult to follow over longer time periods, dynamic CKAP5 dots remained at MT ends after π-EB1 photo-dissociation at time points at which persistent MT growth has completely ceased (Fig. 4; Supplementary Video 5). Thus, EB1-mediated +TIP complexes including endogenous amounts of SLAIN2 do not recruit CKAP5 to MT ends, which might be expected from their distinct non-overlapping binding domains along MT ends^28^. Importantly, however, in cells CKAP5 localization to MT ends by itself is also not sufficient to promote fast, persistent MT growth, but requires an EB1-dependent +TIP activity, which is distinct from the synergistic MT growth acceleration by CKAP5 and EB1 observed *in vitro*^29^.

**Figure 4.**
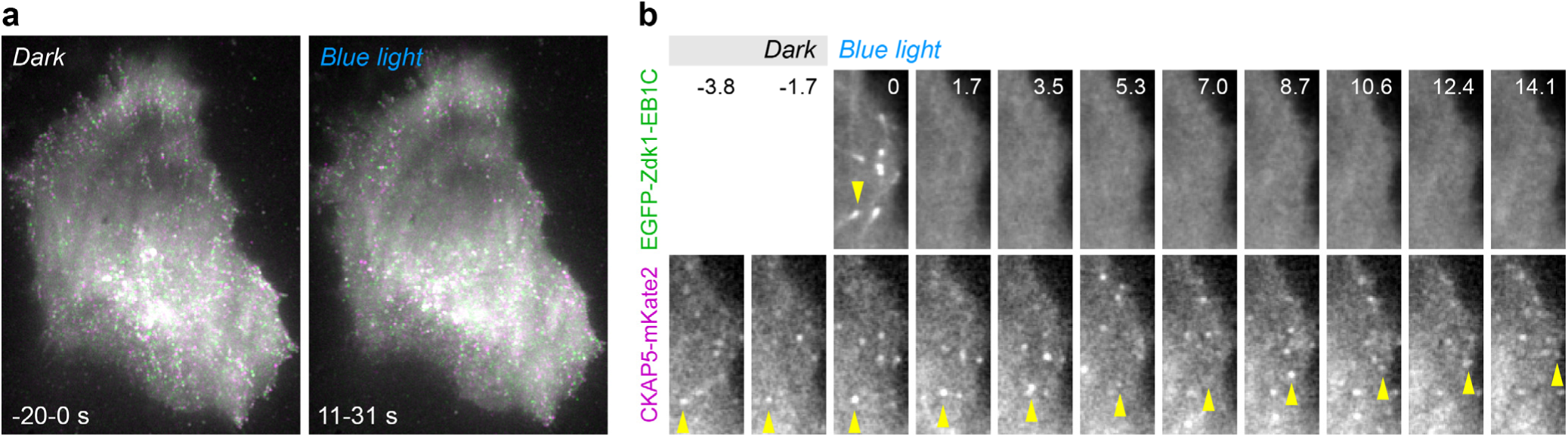
The MT polymerase CKAP5 remains on MT ends upon π-EB1 photo-dissociation. **(a)** Projections of CKAP5-mKate2 TIRF microscopy time-lapse sequences over 20 s temporal windows just prior to and during blue light exposure. Although tracks of MT-end associated CKAP5 dots appear disorganized after π-EB1 photo-dissociation, likely because of attenuated MT growth, they are otherwise similar in appearance and intensity. **(b)**Examples of CKAP5 dot dynamics following π-EB1 photo-dissociation. Arrow highlights CKAP5 that appears to remain associated with a MT end during blue light exposure.

Lastly, we tested if and how acute photo-dissociation of the MT +TIP complex altered intracellular MT organization in tubulin-mCherry expressing π-EB1 cells. Although difficult to quantify precisely, the density of the peripheral MT network dropped rapidly during, and recovered within minutes after terminating blue light exposure (Fig. 5a). An overall decrease of polymerized MTs was further supported by a robust increase in cytoplasmic tubulin-mCherry fluorescence, which reached a new steady state in less than a minute (t_1/2_ = ˜13 s; Fig. 5b) as individual MTs depolymerized in response to π-EB1 photo-dissociation (Fig. 5c; Supplementary Video 6). Interestingly, two distinct MT populations appeared to respond differently. Many radial MTs underwent catastrophes and rapid shortening events within 5-10 s of π-EB1 photo-dissociation (Fig. 5d) consistent with the observed delay in growth rate attenuation (Fig. 3f). In contrast, an underlying population of more curvy MTs was qualitatively more resistant to π-EB1 dissociation indicating that these MTs are stabilized by mechanisms that no longer require a growth-promoting +TIP complex. In migrating cells, the MT cytoskeleton is polarized forward^30^. Because H1299 cells did not directionally migrate as single cells, to test how EB1-recruited +TIP complexes contribute to MT cytoskeleton polarity, we instead expressed constitutively active Rac1(Q61L) in π-EB1 cells, which induces isotropic leading edge protrusion, and exposes fast MT growth toward the cell edge that counter-balances retrograde F-actin flow^31^. Local π-EB1 photo-dissociation resulted in a dramatic retraction of these MTs from Rac1(Q61L)-induced lamellipodia (Fig. 5e; Supplementary Video 7), indicating that EB1-recruited +TIP complexes are specifically required to maintain fast growth of this MT population, in addition to Rac1-mediated inactivation of MT destabilizing activities^32,33^. Importantly, by using patterned blue light exposure intracellular asymmetry of MT network organization could be generated rapidly and maintained for extended periods of time.

**Figure 5.**
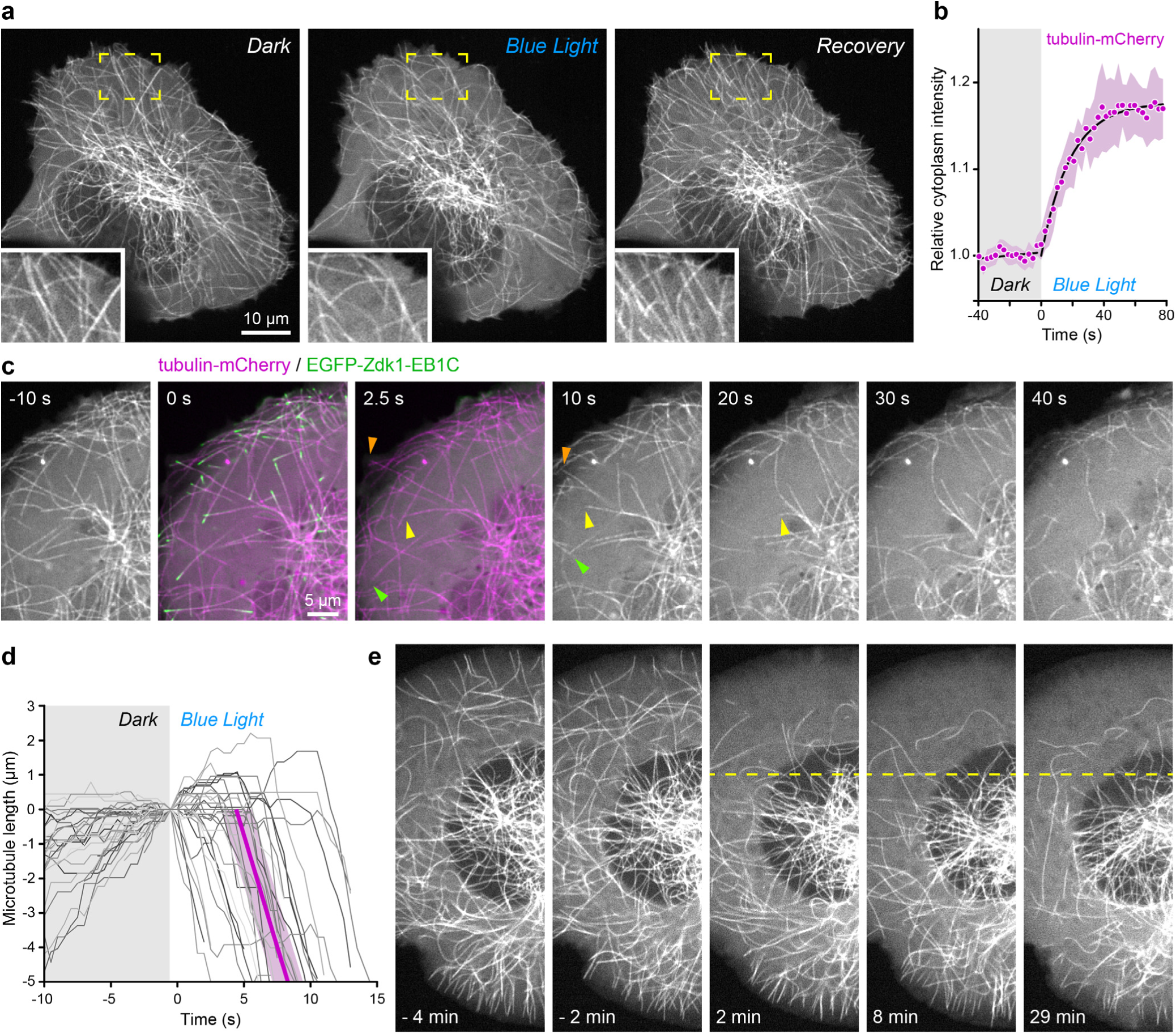
π-EB1 photo-dissociation induced MT cytoskeleton reorganization. **(a)** Tubulin-mCherry expressing π-EB1 cell before, after 30 s during blue light exposure, and after 3 min recovery in the dark. Insets are indicated regions at higher magnification demonstrating reversible decrease of peripheral MT density. **(b)**Analysis of cytoplasmic tubulin-mCherry fluorescence intensity in response to π-EB1 photo-dissociation. n = 5 cells. Shaded area indicates 95% confidence interval. **(c)** Rapid MT depolymerisation in response to π-EB1 photo-dissociation. Arrowheads highlight example MTs. **(d)** Life-history plots of MTs with ends near the cell edge aligned to the time of blue light exposure. n = 33 MTs from 5 cells. The purple line is the average of linear fits of the depolymerizing phase of these MTs. Shaded area indicates 95% confidence interval. **(d)** Time-lapse sequence of tubulin-mCherry in a Rac(Q61L)-expressing π-EB1 cell in response to local blue-light exposure (above the dashed line) illustrating sustained reorganization of the MT network.

In summary, we demonstrate acute and local control of EB1 function in cells by replacing the endogenous protein with a light-sensitive variant. Acute perturbation of the +TIP complex resulted in a new equilibrium of MT polymerization dynamics in less than a minute demonstrating the rapid response time by which cells can reshape their MT network. We find that EB1-mediated +TIP complexes predominantly promote MT polymerization and dominate potential destabilizing effects of direct EB1-binding to MTs, and that unlike *in vitro,* the intrinsic GTP-MT cap is not sufficient to maintain persistent and fast MT polymerization in cells without EB1-recruited +TIP activities. Intriguingly, this +TIP activity does not simply recruit the major MT polymerase CKAP5 to MT ends, but may activate it by other mechanisms that are currently not understood. As a consequence light-sensitive π-EB1 functions as a novel reagent to attenuate MT polymerization and control intracellular MT organization with high spatial and temporal precision, which will be essential to dissect dynamic MT functions in complex cell and tissue morphogenetic processes. Of broader impact, we believe that our approach to insert a light-sensitive protein-protein interaction module between functional domains is a powerful new strategy applicable to a large fraction of the proteome enabling new spatially and temporally resolved experimental questions that cannot be addressed by genetic means alone.

## Methods

### DNA constructs and molecular cloning

mCherry-EB3 was from the Michael Davidson plasmid collection (Addgene plasmid #55037). mCherry-CLASP2γ was cloned by replacing the AgeI/KpnI restriction fragment of pEGFP-CLASP2y^34^ with the corresponding fragment of pmCherrγ-C1. Similarly, mCherry-MCAK was cloned by replacing the AgeI/BsrGI fragment of mEmerald-MCAK (Addgene plasmid #54161, from Michael Davidson) with the corresponding fragment of pmCherry-C1. mCherry-α-tubulin was obtained from Roger Tsien (Addgene plasmid #49149). PCR-based cloning strategies for other constructs are summarized below, and all primers used are listed in supplementary Table 1:

1. EB1N-LZ-LOV2 was constructed as follows: The coding regions corresponding to amino acids 1-185 of an EB1 shRNA-resistant variant^24^ and a GCN4 leucine zipper (generated by oligonucleotide assembly PCR) were first amplified, and then connected by overlap extension PCR, and the resulting product ligated into the NheI/XhoI digested backbone of pEGFP-C1. The LOV2 coding sequence originated from PA-Rac1 (Addgene plasmid #22027)^35^, and L406/407A mutations that were reported to stabilize docking of the Jα-helix^36^ as well as a GSGSG linker sequence were introduced by PCR. This modified LOV2 domain including the linker was amplified and inserted into the SacII/BamHI sites of the EB1N-LZ plasmid.
2. EB1N-LZ-EGFP-LOV2 and EB1N-LZ-mCherry-LOV2 were cloned by inserting PCR amplified EGFP/mCherry coding sequences into XhoI/SacII sites of EB1N-LZ-LOV2.
3. EB1N-EGFP-LZ-LOV2 was cloned by inserting the coding sequence for the PCR-amplified GCN4 leucine zipper into the SacII site of EB1N-GFP-LOV2 by Gibson assembly.
4. mCherry-Zdk1-EB1C was cloned in the following way: PCR-amplified mCherry-Zdk1-linker (from pTriEx-mCherry-Zdk1, Addgene plasmid #81057) and the EB1 C-terminal fragment corresponding to amino acids 186-268 were connected by overlap extension PCR. The resulting PCR product was inserted into the NheI/XhoI sites of pEGFP-C1 .EGFP-Zdk1-EB1C was generated by excising EGFP from pEGFP-C1 by restriction digestion with NheI/BsrGI and ligation into NheI/BsrGI digested mCherry-ZDK1-EB1C, thereby replacing mCherry with EGFP. Untagged ZDK1-EB1 was made by removing mCherry from mCherry-ZDK1-EB1 by restriction digestion with NheI/BsrGI, and removal of overhangs by T4 polymerase, followed by ligation.
5. GST-EB1N-LZ-LOV2 was cloned by inserting PCR amplified EB1N-LZ-LOV2 into the BamHI/XmaI sites of pGEX-4T-2 (GE Life Sciences).
6. 6xHis-Zdk1-EB1C was cloned by inserting PCR-amplified ZDK1-EB1C into NheI/XhoI sites of pET28a.
7. GST-LOV2 was cloned by inserting the PCR-amplified LOV2 coding sequence including a GSGSG linker into BamHI/XhoI sites of pGEX-4T-2.
8. mCherry-Slain2 was cloned by inserting the PCR amplified SLAIN2 coding sequence into the XhoI/EcoRI sites of pBio-mCherry-C1.
9. CKAP5-mKate2 was cloned by inserting the PCR amplified mKate2 coding sequence into the BamHI/NotI sites of chTOG-GFP (Addgene #29480, from Linda Wordeman)

### Cell culture and generation of the π-EB1 cell line

NCI-H1299 human non-small cell lung cancer cells were cultured in RPMI1640 supplemented with 10%FBS, penicillin/streptomycin and non-essential amino acids at 37°C, 5% CO_2_ in a humidified tissue culture incubator. According to NIH guidelines, the identity of NCI-H1299 cells was confirmed by short tandem repeat profiling (IDEXX BioResearch) using a standard 9-marker panel: AMEL(X); CSF1PO(12); D13S317(12); D16S539(12,13); D5S818(11); D7S820(10); THO1(6,9.3); TPOX(8); vWA(16,17,18). The profile obtained was identical to the one reported by the ATCC.

To replace endogenous EB1 with π-EB1 and prevent cells from adapting to EB1 depletion, we first generated stable cell lines expressing both halves of an shRNA resistant variant of π-EB1. H1299 cells were transfected with both untagged EB1N-LOV2 and EGFP-Zdk1-EB1C plasmids and selected with 500 μg/ml G418. Clones expressing both halves at comparable expression levels were identified by microscopy and colonies selected in which EGFP-Zdk1-EB1C was clearly localized to growing MT ends, which requires presence of the unlabelled N-terminal part. Colonies with low MT plus-end to cytoplasm ratio or displaying strong MT lattice labelling were discarded because expression of both halves was expected to be unequal. Expression of both π-EB1 halves was verified by immunoblot using antibodies that specifically recognize either the EB1 N- (Thermo Fisher Scientific Clone 1A11/4) or C-terminal half (BD Biosciences Clone 5/EB1). Next we knocked down endogenous EB1 in these clones by pLKO.1 lentivirus-mediated shRNA. The EB1 shRNA lentivirus and shRNA resistant variants were as described^24^. Selected colonies were sorted at the UCSF Parnassus Flow Cytometry Core to obtain a homogenous population of cells. π-EB1 cells were then transfected with mCherry-tagged constructs for functional experiments either using Fugene6 (Roche) or Lipofectamine 2000 (Life Technologies) according to manufacturer’s instructions.

### EB1/3 immunofluorescence

Cells grown on clean #1.5 glass coverslips (64-0713, Warner Instruments) were fixed in -20°C methanol for 10 min, washed with PBS and blocked with blocking buffer (PBS, 2% BSA, 0.05% Tween 20) for 30 min. Mouse anti-EB1 (BD Biosciences Clone 5/EB1) and rat anti-EB3 (KT36, Absea) antibodies were diluted in blocking buffer (1:150) and 50 μl drops were spotted on a sheet of parafilm in a humidified chamber. Coverslips were placed on top of these droplets (cells facing down), and incubated for one hour at room temperature, after which the coverslips were washed 3 times in PBS, 0.05% Tween 20. Fluorescent secondary antibodies (Alexa488 or Alexa568 conjugated goat anti-mouse or rat antibodies, Invitrogen) were diluted in blocking solution (1:400), and coverslips incubated and washed as above. Coverslips were dehydrated by brief immersion in 70% ethanol followed by 100% ethanol, air dried and mounted in Mowiol mounting medium (0.1 M Tris-HCl pH8.5, 25% glycerol, 10% Mowiol 4-88).

### In vitro binding assays and native PAGE

GST- and 6xHis-tagged proteins were produced in *E.coli* BL21 using standard protocols. Briefly, bacteria were lysed by three freeze thaw cycles in dry-ice in lysis buffer (TBS, 0.5% TX-100, 1mM PMSF, 0.5mg/ml lysozyme) followed by DNAse treatment (1μg/ml). Lysates were cleared by centrifugation at 16000g for 10 min at 4°c, and incubated with glutathione sepharose (GE Healthcare), or Talon resin (Clontech). Affinity resins were washed twice with low salt buffer (TBS, 0.1% TX-100, 5mM 2-mercaptoethanol), three times with high salt buffer (TBS, 650mM NaCl, 0.1% TX-100, 5mM 2-mercaptoethanol). For purification of LOV2-containing constructs, an additional incubation step with low salt buffer containing 5 mg/ml riboflavin 5’- phosphate (FMN) to ensure stoichiometric loading of the LOV2 domain with the FMN co-factor. Proteins were eluted in low salt buffer containing either 25 mM reduced glutathione (pH 8.5), or 150mM imidazole, dialyzed and concentrated (Amicon Ultra-15 centrifugal filters, 3000MWCO), and aliquots snap-frozen in liquid nitrogen.

Interaction between LOV2 and Zdk1 was monitored by discontinuous native PAGE. Purified LOV2 and Zdk1 fusion proteins were mixed and incubated for 30 min at 4°C, and LOV2/Zdk1 complexes run on 6% native PAGE gels in Tris-Glycine buffer pH 8.3 at 75 V for 2 hours. A custom made 470 nm LED array was placed in front of the electrophoresis tank to photoactivate LOV2 as the proteins migrated through the gel.

To test π-EB1 +TIP-binding *in vitro,* H1299 cells were lysed in ice cold 50 mM Tris-HCl, 150 mM NaCl, 0.1% Triton X-100, containing protease inhibitors (cOmplete Protease Inhibitor Cocktail, Sigma) for 30 min. Lysates were cleared by centrifugation at 13000 rpm in an Eppendorf microcentrifuge, and added to glutathione sepharose beads loaded with GST-EB1N-LZ-LOV2 and 6xHis-Zdk1-EB1C, incubated on ice for 30 min, and washed extensively. Bound proteins were analyzed by SDS-PAGE, wet transfer to nitrocellulose (GVS, 1212590) for 1 h at 60V (Bio-Rad Mini Trans-Blot), and chemiluminescent detection with horseradish peroxidase-conjugated secondary antibodies using a Fluorchem Q gel documentation system (92-14116-00, Alpha Innotech) using standard protocols. Primary antibodies: Mouse anti-p150_Glued_ (BD Transduction Laboratories); rat anti CLASP2 (KT68, Absea).

### Microscopy, photoactivation and image analysis

Fluorescent protein dynamics were imaged by spinning disk or TIRF microscopy on a customized microscope setup essentially as described^25,37,38^, except that the system was upgraded with a next generation scientific CCD camera (cMyo, Photometrics) with 4.5 μm pixels allowing optimal spatial sampling using a 60x NA 1.49 objective (CFI APO TIRF; Nikon). Global π-EB1 photo-dissociation was achieved by turning on the 488 nm excitation channel. An irradiance of ˜250 mW/cm^2^ was sufficient to photoactivate the LOV2 domain although simultaneous imaging of EGFP-tagged proteins required higher light intensity. Irradiance at the specimen plane was measured using a X-Cite XR2100 light power meter (EXFO Photonic Solutions).

To achieve subcellular π-EB1 photo-dissociation, a digital micromirror device (Polygon 400, Mightex) equipped with a 470 nm LED was mounted on a Nikon TI auxiliary camera port equipped with a beamsplitter such that 20% of the light path was diverted for photoactivation and 80% was used for spinning disk confocal microscopy. This allowed for direct imaging of the reflected illumination pattern to accurately focus the Polygon 400 as well as simultaneous imaging and π-EB1 photo-dissociation. To eliminate scattered photoactivation light in the imaging channel, the Polygon 400 was operated such that short 10-20 ms blue light pulses were triggered to occur between image acquisitions. For fast time-lapse experiments (at or above 1 frame per second), the Polygon 400 was directly triggered by the camera using a short delay of a few hundred ms after camera exposure. For slower time-lapse experiments, the camera trigger was used to start a pre-programmed pulse sequence between exposures using an ASI MS2000 Sequencer module.

Image analysis was performed in NIS Elements 4.3 or Fiji^39^. Fluorescence intensities were measured in small regions of 3-5 pixel diameter on MT ends that were moved in time with MT growth, and nearby cytoplasm as local background. MT end / cytoplasm ratios were calculated as previously described^40^. A ratio of one indicates no measurable difference between MT end and local cytoplasm. MT plus end tracking of mCherry-EB1N was done using u-track 2.1.3 (available from Gaudenz Danuser at http://www.utsouthwestern.edu/labs/danuser/software/)^41,42^. ‘Comet Detection’ parameters were adapted to decrease the number of false positive detections as follows: ‘High-pass Gaussian standard deviation’: 6; Watershed segmentation ‘Minimum threshold’: between 6 and 8; other parameters remained at default settings. Similarly, in the ‘Tracking’ step, the ‘Minimum Length of Track Segments’ was set to 4. In order to not bias average MT growth rates toward shorter tracks, frame-to-frame MT growth rates were extracted from the ‘tracksFinal’ structure using a custom MatLab script, and frame-to-frame displacements of less than 0.5 pixels were excluded. Tubulin-mCherry labelled MTs ends were tracked using the Fiji MtrackJ plugin^43^.

Statistical analysis was done with the Analyse-It plugin for Microsoft Excel. Statistical significance of multiple comparisons was calculated using the Tukey-Kramer HSD test after confirming normal distribution of the data by Shapiro Wilk testing. Least square curve fitting was performed using the Solver plugin in Microsoft Excel^44^. Figures were assembled in Adobe Illustrator CS5, and videos using QuickTime Pro.

## Author Contributions

J.v.H. and T.W. designed experiments, analysed data and wrote the manuscript. J.v.H. performed most experiments. A.E. performed experiments and generated reagents. H.W. and K.H. contributed unpublished reagents.

## Acknowledgements

This work was supported by National Institutes of Health grants R01 GM079139, R01 GM094819, and S10 RR26758 to T.W. We thank all members of the CTB community for discussions and comments on the manuscript.

## SUPPLEMENTARY DATA

**Supplementary Figure 1.**
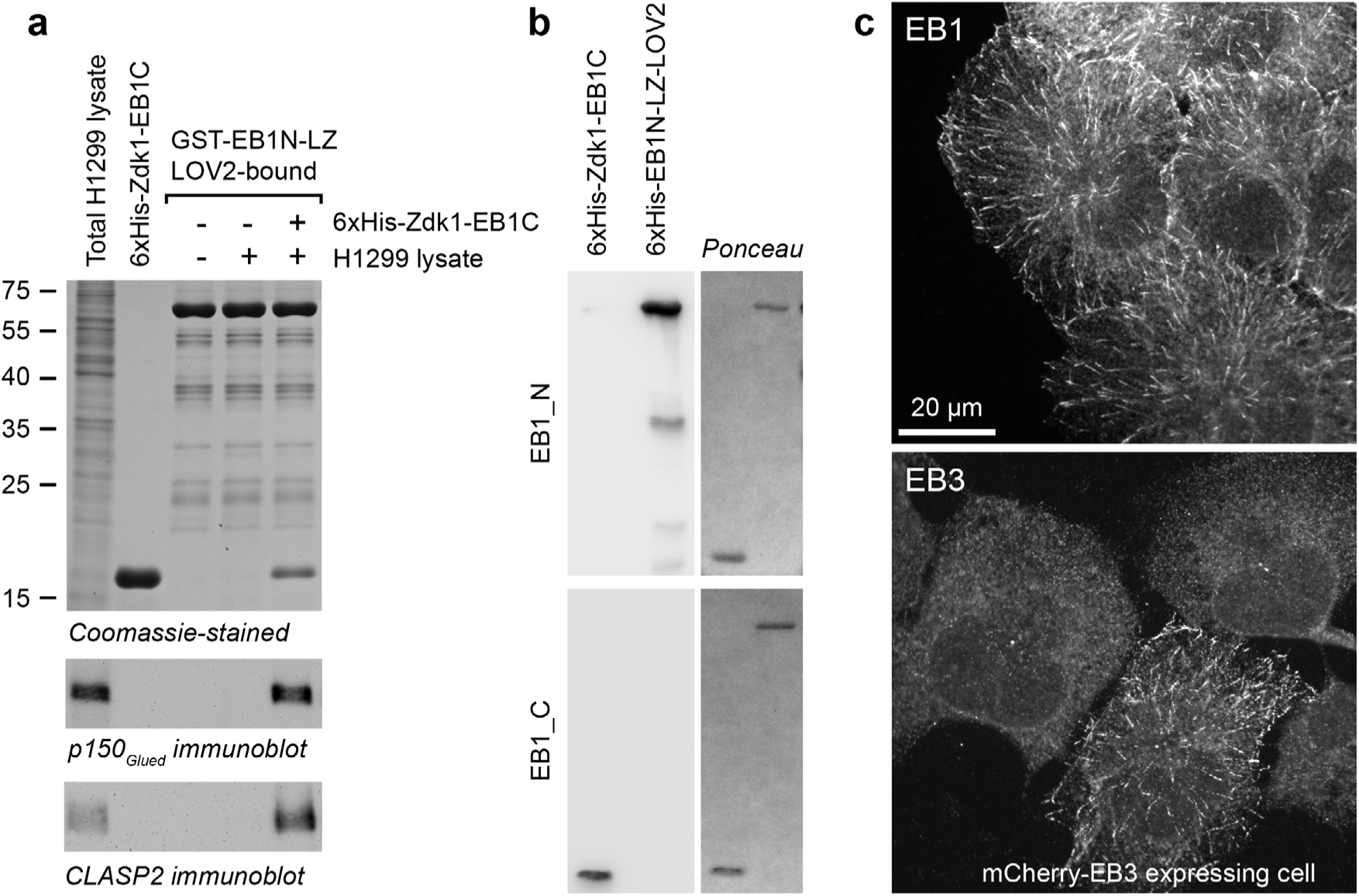
**(a)** GST pull‐down assay of +TIPs from H1299 cell lysate by bacterially produced purified π‐EB1 (GST‐EB1N‐LZ‐LOV2 and 6xHis‐Zdk1‐EB1C). CLASP2 and p150_Glued_ were only isolated from cell lysate in the presence of Zdk1‐EB1C indicating a functional interaction of π‐EB1 with representative +TIPs. **(b)** Immunoblots of the indicated bacterially produced purified 6xHis‐tagged proteins probed with commercially available anti‐EB1 antibodies demonstrating the specificity for either the N‐ or C‐terminal half of π‐EB1. EB1_N: Thermo Fisher Scientific Clone 1A11/4; EB1_C: BD Biosciences Clone 5/EB1. Ponceau staining shows loading of the probed blots. **(c)** H1299 cells express high levels of EB1 but not EB3. Cells were stained as indicated for EB1 (BD Biosciences Clone 5/EB1) or EB3 (Absea Biotechnology KT36). While all cells express high levels of EB1, only cells transiently transfected with mCherry‐EB3 were labelled by the EB3 antibody.

**Supplementary Figure 2.**
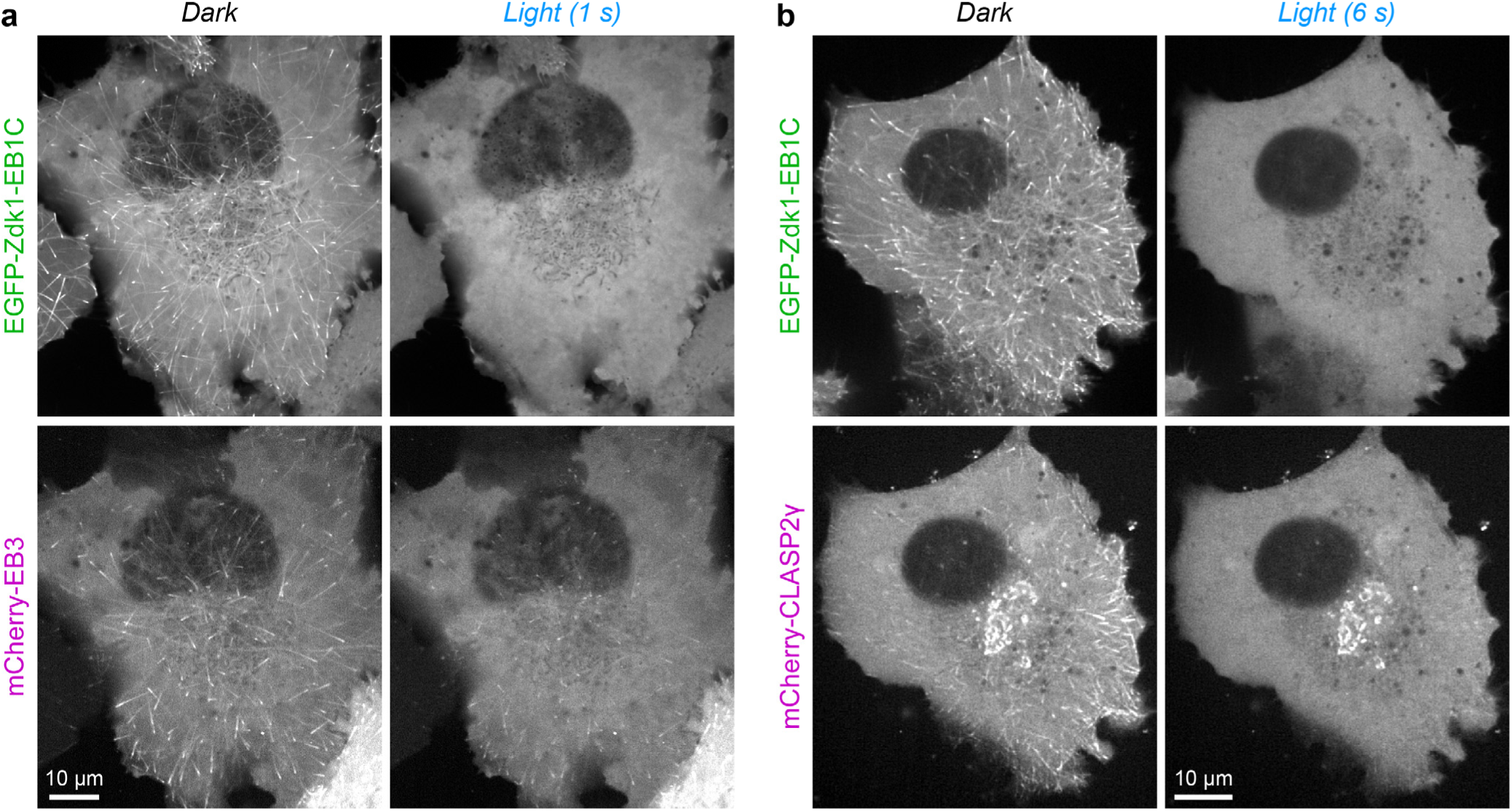
**(a)** π‐EB1 photo‐dissociation partially dissociates transiently transfected mCherry‐EB3 from growing MT ends. This is likely due to heterodimerization of different EBs and indicates that π‐EB1 may act as a dominant negative even in the background of remaining endogenous EBs. **(b)** π‐EB1 photo‐dissociation results in dissociation of transiently transfected mCherry**‐**CLASP2γ from growing MT ends. mCherry**‐**CLASP2γ remains at the Golgi apparatus to which it binds independent of microtubules, indicating specific disruption of MT interactions only. In both**(a)** and **(b)**the dark panel is the first exposure at 488 nm.

**Supplementary Table 1:**
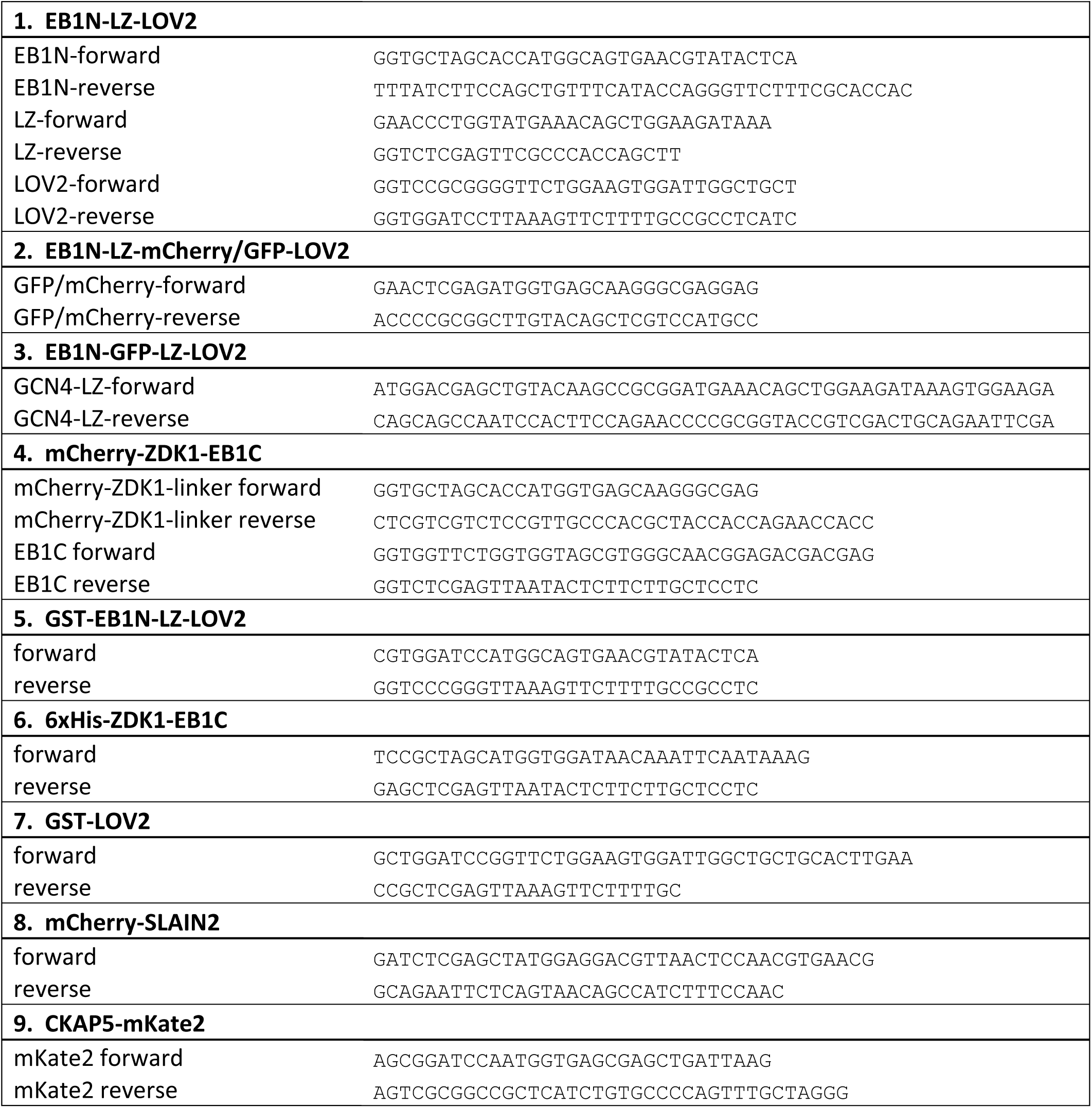
PCR primer sequences used for cloning as described in the methods section.

### Supplementary Videos

**Video 1.** Time-lapse sequence of five consecutive cycles of π-EB1 photo-dissociation showing immediate and reversible release of both the EGFP-tagged π-EB1 C-terminal half and mCherry-SLAIN2 from growing MT ends when 488 nm blue light is turned on. Images were acquired every 2 seconds with a 3 min dark recovery phase between cycles.

**Video 2.** Time-lapse sequence of mCherry-Zdkl-EBlC showing local and reversible π-EBl photodissociation by alternating patterns of blue light exposure. The boundary sharpness in this video illustrates the best we can currently achieve. Images were acquired at one frame per second, and playback is accelerated 15x.

**Video 3.** Time-lapse sequence of EB1N-mCherry-LOV2 before, during and after blue light exposure showing reversible inhibition of MT growth as a result of π-EB1 photo-dissociation. Images were acquired at two frames per second, and playback is accelerated 30x.

**Video 4.** Time-lapse sequence of EB1N-mCherry-LOV2 showing local inhibition of MT growth. Only the top half of the cell is exposed to 470 nm blue light during the 40-79 second time window. Images were acquired at one frame per second, and playback is accelerated 15x.

**Video 5.** Time-lapse sequence of CKAP5-mKate2 dynamics by TIRF microcopy before and during of π-EB1 photo-dissociation. Dynamic CKAP5-mKate2 dots remain during blue light exposure that indicating that CKAP5 association with the most distal tip of growing MTs does not depend on functional EB1. Top panel shows the EGFP-tagged C-terminal half of π-EB1 dissociating from MT ends when 488 nm exposure is turned on.

**Video 6.** Time-lapse sequence of tubulin-mCherry before and during blue light exposure illustrating rapid depolymerization of a population of microtubules in response to π-EB1 photo-dissociation. Left panel shows the EGFP-tagged C-terminal half of π-EB1 dissociating from MT ends when 488 nm exposure is turned on. Images were acquired every 2.5 seconds and playback is accelerated 6x.

**Video 7.** Time-lapse sequence of tubulin-mCherry in a Racl(Q6lL)-expressing π-EB1 cell. Only approximately the right third of the cell is exposed to blue light resulting in sustained retraction of the MT network from this part of the cell. Images were acquired every 30 seconds and playback is accelerated 450x.

